# Activated thiol Sepharose-based proteomic approach to globally quantify protein oxidation

**DOI:** 10.1101/549709

**Authors:** Yang Xu, Benjamin G Neel

**Affiliations:** Department of Medical Biophysics, University of Toronto, Toronto, Ontario, Canada M5G 1L7; Princess Margaret Cancer Centre, University Health Network, Toronto, Ontario, Canada M5G 1L7; Laura and Isaac Perlmutter Cancer Center, New York University Langone Medical Center, New York, New York 10016, USA

**Author notes:** To whom correspondence should be addressed: Benjamin G. Neel: Laura and Isaac Perlmutter Cancer Center, New York University Langone Medical Center, New York, New York 10016, USA.

**Keywords:** protein oxidation, redox regulation, PKM2, HLRCC, ATSP, proteomics

## Abstract

Reactive oxygen species (ROS) can act as second messengers for various signaling pathways, and abnormal oxidation contributes to multiple diseases, including cancer. Detecting and quantifying protein oxidation is crucial for a detailed understanding of redox signaling. We developed an Activated Thiol Sepharose (ATS)-based proteomic approach (ATSP) to globally quantify protein oxidation. ATSP can enrich for H_2_O_2_-sensitive thiol peptides, which are more likely to contain reactive cysteines involved in redox signaling. We applied our approach to analyze hereditary leiomyomatosis and renal cell carcinoma (HLRCC), a type of kidney cancer that harbors *fumarate hydratase* (*FH*)-inactivating mutations and has elevated ROS levels. Multiple proteins were oxidized in FH-deficient cells, including many metabolic proteins, such as the M2 isoform of pyruvate kinase (PKM2). Treatment of HLRCC cells with dimethyl fumarate (DMF) or PKM2 activators alter PKM2 oxidation levels. This newly developed redox proteomics method can globally detect oxidation of reactive cysteines and can be employed to analyze multiple physiological and pathological conditions.

## Introduction

Over the past two decades, increasing evidence has revealed that oxidation of cysteinyl residues is an important post translational modification (PTM) that can regulate enzymatic activity, protein localization, and/or protein-protein interactions (1–3). However, compared with other PTMs, protein oxidation is understudied, at least in part because of technical difficulties in detecting oxidation events (4), such as the instability of some protein oxidation states, post-lysis oxidation from ambient oxidants (mainly O_2_), signal attenuation from surrounding antioxidants, and low signal to noise ratio due to the transience of protein oxidation during signal transduction.

Multiple techniques attempt to overcome these difficulties. Some were designed to specifically detect oxidation of protein tyrosine phosphatases (PTPs), which are particularly sensitive to oxidation. The modified in-gel phosphatase assay was the first to detect oxidation of multiple PTPs simultaneously (5,6). Subsequently, our group utilized a specific antibody against the most highly oxidized state of classical PTPs to develop a mass spectrometry (MS)-based approach to globally detect their oxidation (7,8). However, many other types of protein can be oxidized, and in several cases, such oxidation has important physiological and pathological consequences (9–12). More general redox techniques have been developed to globally detect protein oxidation, such as the isotope-coded affinity tag (ICAT) method (13), the modified cysteinyl-labeling assay (14,15), and dimedone/dimedone antibody-based approaches (16–19). Nevertheless, each has certain technical limitations. For example, the ICAT method measures decreases in oxidation signal after cellular stimulation, resulting in an intrinsically low signal to noise ratio (4). Although dimedone analogs directly label oxidized cysteines, they mainly react with sulfenic acids (S-OH), which are only one of the oxidation states of cysteine. Because the sulfenic acid state is usually labile and rearranges rapidly to a sulfenylamide (S-N) or disulfide (S-S) (20–22), dimedone/dimedone antibody-based approaches might miss important oxidation events, especially in cells under chronic oxidative stress, an important feature of many pathologies.

Activated Thiol Sepharose (ATS), which contains a dipyridyl disulfide group that can bind to free thiol-containing proteins (23), is another promising redox probe. The disulfide bond formed between ATS and free thiols is both strong and reversible, enabling harsh purification conditions, followed by easy, quantitative elution for MS analysis. Several groups have used ATS to assess cysteine oxidation or other cysteine modifications inside cells (24–27). Those studies all featured a protein denaturation step before ATS labelling, which disrupts the local environment of the reactive cysteines (involved in redox signaling), rendering them indistinguishable from structural cysteines (not involved in redox signaling).

We developed an ATS-based proteomic approach (ATSP) that preserves, and thus enriches for, reactive cysteines. We applied ATSP to *fumarate hydratase* (*FH*)-mutated HLRCC cell lines and identified multiple proteins, especially metabolic proteins, that were oxidized in these cells, including PKM2 and transketolase (TKT). PKM2 oxidation was confirmed by immunoblot (IB) and could be modulated by treating cells with DMF or PKM2 activators. We conclude that ATSP has great potential for detecting reactive cysteine oxidation in redox signaling.

## Results

### Development of ATSP to detect protein oxidation

Reactive cysteines usually have lower pK_a_s than typical cysteines (pK_a_~8.5). Consequently, reactive cysteines exist as thiolate anions (S^−^) at physiological pH, rendering them more chemically reactive to substrates, ROS, and/or other chemicals (28). For ATSP, cells are lysed in de-gassed, non-denaturing lysis buffer containing *N*-ethylmaleimide (NEM); under these conditions, reduced (but not oxidized) cysteinyl residues react with the NEM. Oxidized (reactive) cysteinyl residues that have escaped alkylation are then reduced with dithiothreitol (DTT), incubated with ATS resin, placed in sample buffer for immunoblotting, or processed for MS (Fig. 1A and Methods). To enhance the specificity of labeling, the ATS incubation is performed at acidic pH with a limited amount of ATS resin (29,30) (also, see below). For MS analysis, ATS-bound proteins are washed in denaturing wash buffer, subjected to two-step on-resin trypsin digestion, eluted with tris(2-carboxyethyl)phosphine (TCEP), and alkylated by iodoacetamide (IAM). ATSP can be modified to quantify total protein levels (ATSP-Total) by leaving out the initial NEM alkylation step and using a sufficient amount of ATS resin (Fig. 1A and Methods). The minimal amount of ATS resin for ATSP is determined empirically for each cell system. For example, as shown in Fig. 1B, a given amount of Rat1 cell lysate was processed by ATSP-Total, with sequentially increased amounts of ATS resin. As the readout, we used SHP2, which, as a member of PTP superfamily, has a very reactive catalytic cysteinyl residue. The lowest amount of ATS (20 μL in this case) that can deplete SHP2 from supernatant is used for ATSP (Fig. 1B).

**Figure 1.**
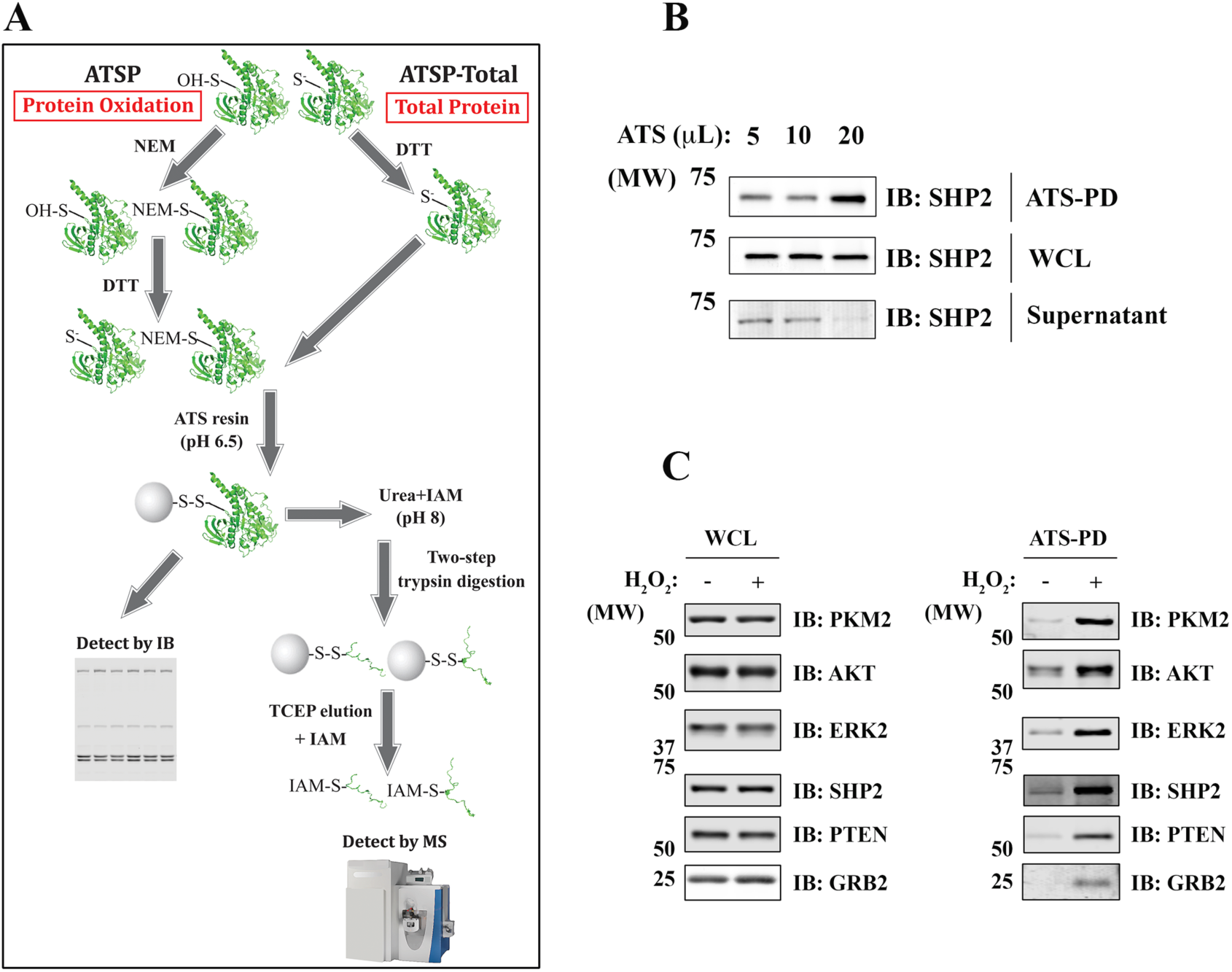
Development of ATSP to detect protein oxidation. A, schematic of ATSP. To detect protein oxidation (ATSP), reduced cysteinyl residues are sequestered by NEM, while the oxidized cysteinyl residues are reduced by DTT before addition of ATS. ATS-bound proteins are subjected to sequential on-resin trypsin digestion, eluted with TCEP, alkylated by IAM, and analyzed by MS. For detection of total proteins (ATSP-Total), all procedures are the same except for the initial alkylation step. B, determination of the amount of ATS resin used for ATSP. Rat1 cells were processed by ATSP-Total, and whole cell lysate (WCL), ATS-bound proteins (pull down, PD), and the supernatant were immunoblotted for SHP2. C, oxidized proteins detected by ATSP. Rat1 cells were treated with 1 mM H_2_O_2_ (4 min) or left untreated. After processing for ATSP, WCL and the ATS-bound proteins were analyzed by IB.

We used ATSP, followed by immunoblotting, to assess protein oxidation in Rat1 cells treated with H_2_O_2_ (Fig. 1C). Multiple previously reported ROS-sensitive proteins were oxidized upon H_2_O_2_ treatment, including PTPs (SHP2 and PTEN), kinases (AKT and ERK2), an adaptor protein (GRB2), and a glycolytic enzyme (PKM2). We also processed the same H_2_O_2_-treated samples by MS. Approximately 3,300 unique cysteine-containing peptides, comprising ~1,300 distinct proteins were detected by MS (Table 1A). As expected, ATSP highly enriched for cysteine-containing peptides, with more than 98% of all the peptides identified by MS containing a thiol group (Table 1A). In addition, of the ~3,300 cysteine-containing peptides, ~1,800 (representing ~1,000 proteins) showed significantly increased oxidation after H_2_O_2_ treatment; these proteins are likely to contain low pK_a_ reactive cysteines (Supporting information).

**Table 1.** Total number of peptides and number of IAM-modified peptides by ATSP. A, Rat1 cells were treated with 1 mM H_2_O_2_ or left untreated. After processing for ATSP and analyzing with MS, the number of total peptides and the cysteine-containing peptides (IAM-modified) were listed (mean ± SEM; n=3). B, UOK262 FH-WT, FH-deficient, and FH-WT-H_2_O_2_ (1 mM; 4 min) were processing for ATSP and analyzed by MS. The number of total peptides and the cysteine-containing peptides (IAM-modified) were listed (mean ± SEM; n=3).

| **A** |  |  |  |
| --- | --- | --- | --- |
|  | IAM-modified peptides | Total peptides | Percentage of IAM-modified peptides |
| No treatment | 2479 ± 88 | 2517 ± 90 | 98.48% ± 0.24% |
| H<sub>2</sub>O<sub>2</sub> | 3283 ± 189 | 3340 ± 188 | 98.27% ± 0.12% |

| **B** |  |  |  |
| --- | --- | --- | --- |
|  | IAM-modified peptides | Total peptides | Percentage of IAM-modified peptides |
| FH-WT | 3416 ± 165 | 3535 ± 162 | 96.64% ± 0.30% |
| FH-deficient | 3380 ± 275 | 3522 ± 289 | 95.97% ± 0.38% |
| H<sub>2</sub>O<sub>2</sub> | 4182 ± 16 | 4326 ± 25 | 96.66% ± 0.19% |

### Metabolic proteins were highly oxidized in FH-deficient HLRCC cells

HLRCC is an inherited kidney cancer syndrome, associated with inactivating mutations in *fumarate hydratase* (*FH*), which encodes a TCA cycle enzyme (31). FH loss-of-function leads to accumulation of fumarate, resulting in increased ROS levels (32,33). To assess protein oxidation in HLRCC cells, we applied ATSP to the patient-derived HLRCC cell line UOK262 (FH-deficient), compared with cells in which wild-type *FH* had be re-introduced (UOK262 FH-WT) (34). Similar to the results seen in H_2_O_2_-treated Rat1 cells, ~96% of the MS-recovered peptides from UOK262 cells contained cysteinyl residues (Table 1B). Furthermore, multiple proteins were highly oxidized in FH-deficient cells, including many metabolic enzymes (Fig. 2A), antioxidant proteins (Fig. 2B), and various other proteins, such as those involved in the cytoskeleton, translation and chaperones (Fig. 2C). Among the oxidized metabolic enzymes, multiple cysteines on PKM2 (from the glycolysis pathway), TKT (from the pentose phosphate pathway/PPP), and PSAT1 (from serine synthesis pathway/SSP) had the highest oxidation levels in FH-deficient cells (Fig. 2A).

**Figure 2.**
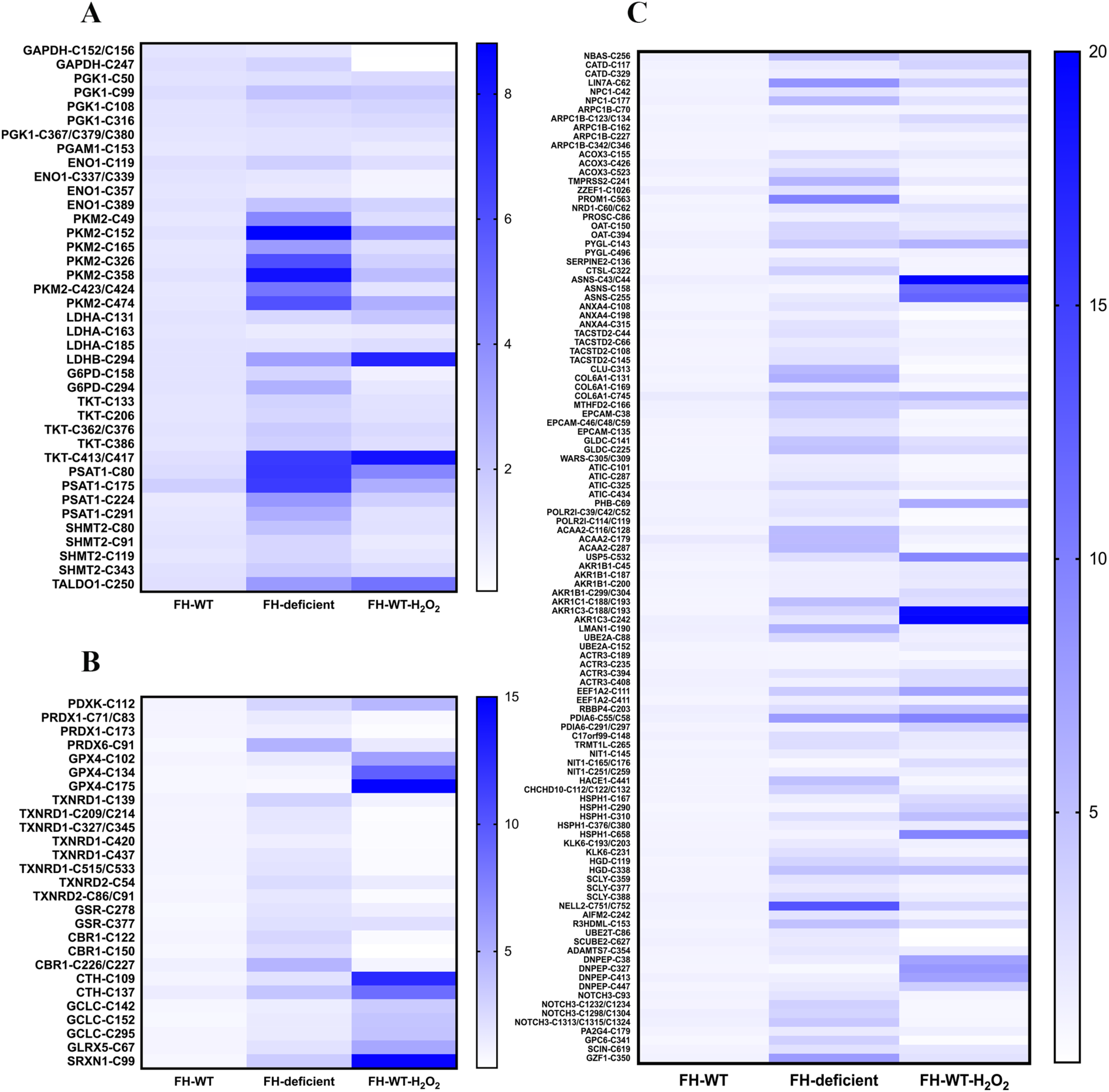
Protein oxidation in FH-deficient HLRCC cells, detected by ATSP-MS. UOK262 (FH-WT or FH-deficient) cells were processed for ATSP. Metabolic proteins are shown in (A), antioxidant proteins are shown in (B), and other proteins are shown in (C). Data were analyzed by label-free quantification of peptides from each identified protein (n=2-3). H_2_O_2_, 1 mM for 4 min.

We used ATSP-IB to validate some of the ATSP-MS results. As predicted by ATSP-MS, PKM2 and TKT were oxidized in FH-deficient, compared with FH-WT, cells (Fig. 3A). The total amount of G6PD and PHGDH, which are involved in the PPP and SSP, respectively, were increased significantly in FH-deficient cells. By contrast, TKT and PKM2 showed only a little or no increase in expression in FH-deficient cells (Fig. 3A). Similar results were obtained by using another HLRCC patient-derived cell line, UOK268, again comparing FH-deficient cells with FH-WT cells (Fig. 3B). To assess the stoichiometry of PKM2 oxidation, we compared the amount of oxidized PKM2 (using ATSP) with the total amount of PKM2 (using ATSP-Total). On average ~20-30% of PKM2 was oxidized in UOK262 FH-deficient cells (Fig. 3C).

**Figure 3.**
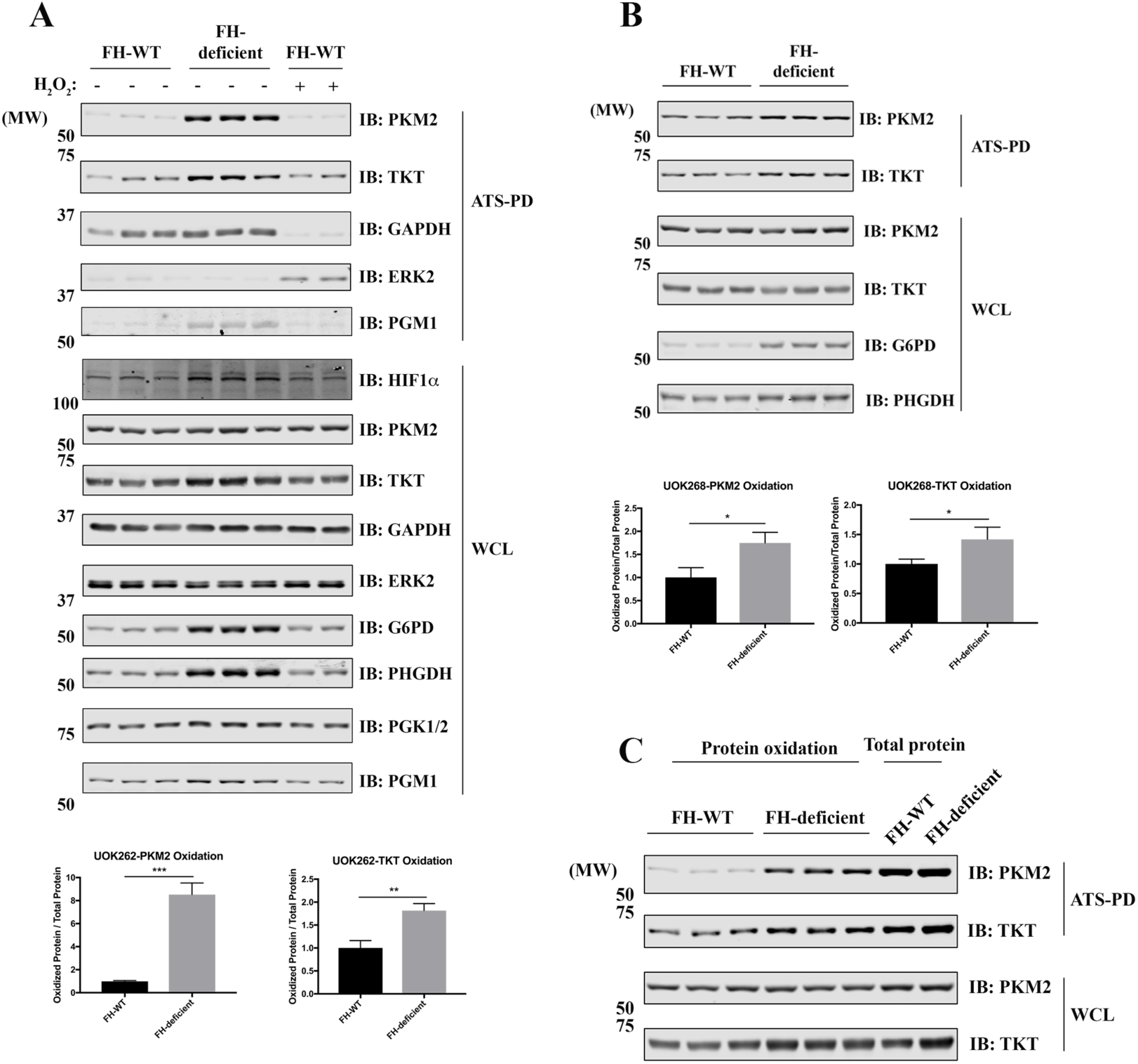
PKM2 is oxidized in FH-deficient cells. A and B, UOK262 (A) and UOK268 (B) (FH-WT and FH-deficient) cells were processed for ATSP. WCL and ATS-bound proteins were analyzed by immunoblotting with the indicated antibodies. The relative amounts of PKM2 and TKT oxidation are quantified below. Data represent mean ± SEM (n=6; *p<0.05, **p<0.01, ***p<0.001; unpaired two-tailed t-test). C, UOK262 (FH-WT and FH-deficient) cells were processed for ATSP or ATSP-Total. WCL and ATS-bound proteins were analyzed by immunoblotting with the indicated antibodies.

### DMF and PKM2 activators can modulate PKM2 oxidation

To test whether PKM2 oxidation is related to intracellular accumulation of fumarate, we treated UOK262 FH-WT cells with dimethyl fumarate (DMF), a cell-permeable fumarate analog. Long-term DMF treatment of FH-WT cells induced PKM2 oxidation and G6PD and PHGDH upregulation to levels similar to those in FH-deficient cells, indicating that the observed protein oxidation and/or overexpression is likely to be the direct consequence of accumulation of fumarate in FH-deficient cells (Fig. 4A).

**Figure 4.**
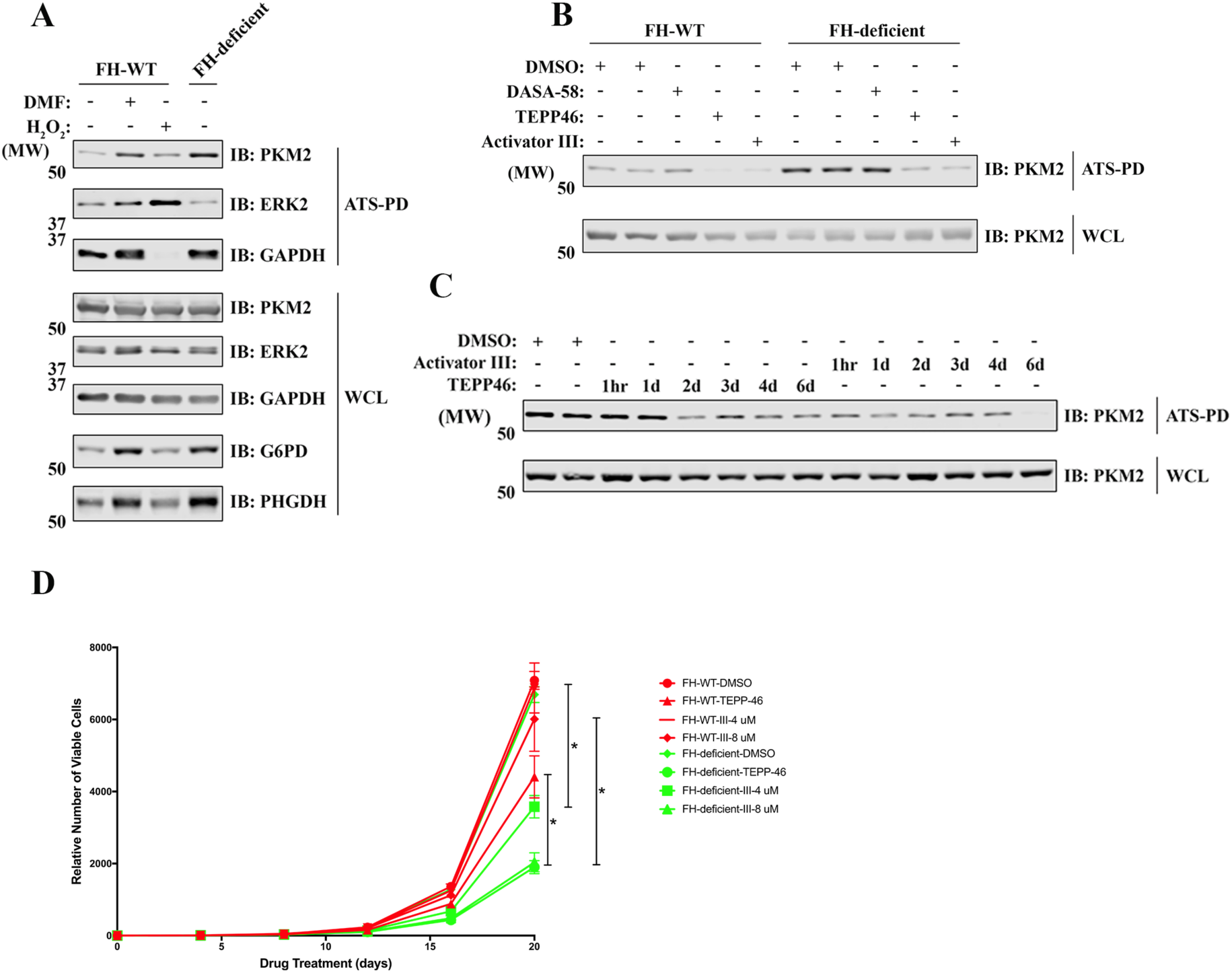
DMF and PKM2 activators can modulate PKM2 oxidation. A, UOK262 cells were treated with DMF (100 μM, 7 days) or H_2_O_2_ (1 mM, 4 min) and processed for ATSP. WCL and ATS-bound proteins were analyzed by immunoblotting with indicated antibodies. B, UOK262 cells were treated with various PKM2 activators (DASA58, 20 μM for 7 days; TEPP46, 20 μM for 7 days; Activator III, 8 μM for 7 days), and processed for ATSP. WCL and ATS-bound proteins were analyzed by immunoblotting with anti-PKM2 antibody. C, FH-deficient UOK262 cells were treated with PKM2 activators (20 μM TEPP46 or 8 μM Activator III) for the indicated time points and processed for ATSP. WCL and ATS-bound proteins were analyzed by immunoblotting with anti-PKM2 antibody. D, UOK262 (FH-WT and FH-deficient) cells were treated with PKM2 activators or DMSO control for the indicated time points. The relative number of viable cells was counted after drug treatments. Data represent mean ± SEM. (n=3; *p<0.05, **p<0.01, ***p<0.001; unpaired two-tailed t test).

Oxidation of PKM2 on C358 has been reported in diamide-treated lung cancer cell lines; moreover, C358 oxidation inhibits tetramer formation and impairs its glycolytic activity (35). Pre-treatment with PKM2 activators, which can promote tetramer formation, can protect reduced PKM2 from diamide-induced oxidation (35). At least seven different cysteine sites on PKM2 were oxidized in HLRCC cells, including C358 (Fig. 2A), so we asked whether different PKM2 activators they can reverse PKM2 oxidation. After a 7-day incubation, two out three PKM2 activators, TEPP46 and Activator III, significantly reduced PKM2 oxidation in UOK262 cells, and both PKM2 activators reduced PKM2 oxidation levels in FH-deficient cells to that seen in FH-WT cells (Fig. 4B). These findings suggest that most oxidized cysteine sites on PKM2 are directly or indirectly involved in PKM2 tetramer formation. Although it took TEPP46 1-2 days to reduce PKM2 in FH-deficient cells, Activator III can reverse PKM2 oxidation within 1-hour of treatment (Fig. 4C), indicating that the redox state of PKM2 is quite dynamic.

Finally, we asked whether preventing/reversing PKM2 oxidation by activator treatment would affect cell viability. Both PKM2 activators had only mild differential growth inhibitory effects on FH-deficient, compared with FH-WT, cells, and this effect was only seen after long term PKM2 activator treatment (Fig. 4D). These results indicate that PKM2 oxidation plays a minor role in cell proliferation of FH-deficient cell, at least under these in vitro conditions.

## Discussion

Redox signaling is involved in a variety of signaling pathways and diseases, and understanding its roles requires sensitive and specific techniques to detect oxidized proteins. We developed ATSP to globally monitor and quantify oxidation of functional thiol-containing proteins. Because reactive cysteines are only a very small subset of all cysteinyl residues and lack canonical consensus sequence motifs, it is quite difficult to directly identify them and study their regulation (30). ATSP capitalizes on the low-pK_a_ of such cysteines (36,37) by: 1) lysing cells in non-denaturing conditions to preserve the structures that confer the low pK_a_s of these reactive cysteines; 2) use relatively acidic labeling buffer to reduce the labeling of non-reactive cysteines (5,29); 3) minimize the concentration of redox probes to label reactive cysteines, whose labeling is less sensitive to the redox probe concentrations than that of non-reactive cysteines (30). In order to get the “limited amount of ATS” for ATSP, we applied ATSP-Total and used SHP2 as a readout. This “limit the amount of ATS” is likely to be adequate for most reactive cysteines because SHP2 depletion was performed on the “total protein” samples while we use it to label the oxidized proteins (typically < 10% of total protein).

We applied ATSP to H_2_O_2_-treated Rat1 cells and found more than half of all the detected peptides showed significant increase in oxidation. Because H_2_O_2_ is a relatively mild ROS and mainly reactive cysteines are ROS-sensitive under low to mild oxidative stresses, ATSP can enrich and specifically characterize the redox states of functional reactive cysteines. By applying ATSP to *FH* mutated HLRCC cell lines, we found multiple metabolic proteins were highly oxidized, including at least 7 cysteine sites on PKM2. Although only C358 has been reported to regulate PKM2 tetramer formation, other cysteine sites might be also directly or indirectly involved in tetramer formation because PKM2 activators can fully reverse PKM2 oxidation. Similar results have been reported for skeletal-type ryanodine receptor (RyR1). Although RyR1-C3635 is selectively nitrosylated, 6 additional cysteines on RyR1 may also contribute to the redox regulation of the channel (38,39). In addition, PKM2 plays important roles in multiple cancers (40), and PKM2 oxidation has been implicated to protect lung cancer cells from oxidative stresses (35). Perhaps surprisingly, then, treatment with either of two PKM2 activators had little differential effect on the proliferation of FH-deficient, compared with FH-WT, HLRCC cells. These results do not, however, exclude the possibility that PKM2 oxidation plays a more important role in the proliferation and/or survival of such cells under the more challenging environment of a tumor. Further studies are needed to evaluate this possibility, as well as to determine what other pathways rescue the effects of PKM2 reactivation *in vitro*.

One potential drawback of ATSP is that the ATS might not “fit” into all molecules with reactive cysteines. We used ATS-4B in all of the above experiments, but we have also tested another commercially available ATS, ATS-6B. ATS-6B works well when using the ATSP protocol to detect proteins oxidation by IB (data not shown), but the difference between ATS-6B and ATS-4B in terms of MS-detected ATS-bound protein profile warrants future investigation. Furthermore, some ATS-bound proteins can be detected by IB, but not MS. This indicates that additional procedures are needed to increase the sensitivity of MS detection, for example, sample fractionation or using a different protease (some tryptic peptides might not fly well in MS instruments).

To summarize, we developed a novel redox approach, ATSP, to globally detect protein oxidation. ATSP is suitable to detect oxidation of reactive cysteines, which are more likely to involved in redox signaling. We applied ATSP to HLRCC cells that have high ROS levels, and we found multiple metabolic proteins were oxidized in these cells, including PKM2. Therefore, ATSP has great potential to be applied in other ROS-related physiological and pathological conditions and help to better understand the redox signaling.

## Materials and methods

### Cell lines and cell culture

Rat1 cells were kindly provided by Dr. Nicholas Tonks (CSH Laboratory, USA). UOK262 FH-WT, UOK262 FH-deficient, UOK268 FH-WT and UOK268 FH-deficient cells were kindly provided by Dr. W. Marston Linehan (NCI, USA). All the cells were cultured in Dulbecco’s modified Eagle’s medium (DMEM). All media contained 10% FBS (Corning) and 100 Units/ml penicillin/streptomycin (Gibco).

### ATS-based proteomic approach

A lysis buffer containing 20 mM Tris-HCl (pH 7.35), 150 mM NaCl, 10% glycerol, 1% NP40 was degassed overnight with vacuum at RT, and then transferred to a hypoxia workstation. NEM (30 mM, Sigma), 0.5 U/μL catalase (Calbiochem), 0.125 U/μL SOD (Sigma) and a protease inhibitor cocktail (2 μg/mL antipain, 2 μg/mL pepstatin A, 20 μg/mL leupeptin, and 20 μg/mL aprotinin) were added to the lysis buffer immediately before using. Within the hypoxia workstation, cells were quickly washed once with degassed PBS, immediately lysed, and incubated for 1 hr at 4°C in the dark with constant rotation. Homogenates were clarified at 10,000× g for 1 min at 4°C. Supernatants were collected, and the buffer was exchanged by gel filtration (GE Healthcare Life Sciences) into the degassed solution (20 mM HEPES [pH 6.5], 100 mM NaCl, 10% glycerol, 0.05% NP40, 0.02 U/μL catalase). Samples were then treated with 1.5 mM DTT and rotated in the dark for 2 hrs at 37°C. Within the hypoxia workstation, lysates were buffer-exchanged again to remove DTT. Activated Thiol Sepharose 4B (GE Healthcare Life Sciences) was added, and the samples were rotated overnight at 4°C in the dark.

For IB, ATS-bound proteins were washed 3× with RIPA buffer, 1× with 8 M urea, and then incubated with 2× Laemmli sample buffer (30 min at 56°C) before resolving on SDS-PAGE. For MS, ATS-bound proteins were first washed 3× with RIPA buffer. ATS-bound proteins were then incubated in a buffer containing 8 M urea and 20 mM IAM (dissolved in 50 mM ammonium bicarbonate [pH 8-8.5]) for 10 min at RT in dark with constant rotation (repeat once). After alkylation, ATS-bound proteins were washed with 50 mM ammonium bicarbonate (pH 8-8.5) (4×), and then digested with 1 μg of trypsin (Thermo Scientific) for 30 min at 37°C with constant rotation. The supernatant after first trypsin digestion was discarded. ATS-bound proteins/peptides were washed once with 50 mM ammonium bicarbonate, and again digested with 1 μg of trypsin overnight at 37°C with constant rotation. The next day, ATS-bound peptides were washed with 50 mM ammonium bicarbonate (3×) and eluted with 1 mM TCEP (Thermo Scientific) for 30 min at 37°C with constant rotation. The eluates were then alkylated with 10 mM IAM (30 min at RT in the dark). The elution/alkylation procedure was repeated one more time, and the two eluates were combined, acidified with trifluoroacetic acid (final pH ~2), purified using StageTips and analyzed by MS.

### Immunoblotting

Whole cell lysate (WCL; 10-50 μg), ATS-bound proteins and/or supernatant were resolved by SDS-PAGE and analyzed by immunoblotting. Antibodies included: anti-PGK1/2, anti-FH, and anti-ERK2 antibodies from Santa Cruz Biotechnology; anti-GAPDH antibody from EMD Millipore; anti-HIF1α antibody from BD Transduction Laboratories; anti-PGM1 and anti-TKT antibodies from Abcam; anti-PHGDH, anti-G6PD, anti-PKM2 antibodies from Cell Signaling Technology.

### LC-MS/MS

The digested sample was loaded onto an Acclaim PepMap trap column in line with an EASY-Spray PepMap analytical column by using the auto sampler of either the EASY-nLC 1000 or EASY-nLC 1200 HPLC with solvent A, consisting of 2% acetonitrile in 0.5% acetic acid, and solvent B, consisting of 80% acetonitrile in 0.5% acetic acid. Peptides were gradient-eluted into either the Orbitrap Elite or Orbitrap Fusion Lumos Mass Spectrometer. MS1 scans were acquired in the Orbitrap and MS2 scans in the ion trap using a data-dependent acquisition scheme. All acquired MS2 spectra were searched against the UniProt reference proteome database using Sequest within Proteome Discoverer. The search parameters were as follows: precursor mass tolerance ± 10 ppm, fragment mass tolerance ± 0.4 Da, trypsin cleavage with 2 missed cleavages allowed, variable modification of oxidation of methionine, carbamidomethyl and N-ethylmaleimide modification of cysteine, and deamidation of glutamine and asparagine. Label-free quantification was obtained by using Skyline (MacCoss Lab Software; version 4.1.0.11796).

### Statistical analysis

Data are presented as the mean ± SEM. Statistical significance was determined using unpaired Student’s t-test. Statistical analyses were performed using Graph-Pad Prism 7, with p < 0.05 considered significant.

## Acknowledgments

We thank Dr. Nicholas Tonks (CSH Laboratory) for providing Rat1 cells, Dr. W. Marston Linehan (NCI) for providing HLRCC cells, and Dr. Beatrix Ueberheide and Mr. Joshua Andrade (PCC Proteomics Shared Resource) for their help with MS. This work was supported by R01CA49152 (BGN) and the Princess Margaret Cancer Foundation.

## The abbreviations used are

ATS: activated thiol Sepharose
ATSP: ATS-based proteomic approach
ROS: reactive oxygen species
HLRCC: hereditary leiomyomatosis and renal cell carcinoma
FH: fumarate hydratase
PKM2: pyruvate kinase M2 isoform
DMF: dimethyl fumarate
PTM: post translational modification
PTP: protein tyrosine phosphatase
MS: mass spectrometry
ICAT: isotope-coded affinity tag
TKT: transketolase
RT: room temperature
IB: immunoblot
WCL: whole cell lysate
NEM: *N*-ethylmaleimide
DTT: dithiothreitol
TCEP: tris(2-carboxyethyl)phosphine
IAM: iodoacetamide
WT: wild-type
PPP: pentose phosphate pathway
SSP: serine synthesis pathway
RyR1: skeletal-type ryanodine receptor

## Notes

**Financial support:** This work was supported by R01CA49152 (BGN) and the Princess Margaret Cancer Foundation.

**Conflicts of interest:** BGN is a co-founder, has equity in, and receives consulting funds from, Northern Biologics, and is a co-founder, chair of the Scientific Advisory Board, has equity in, and receives consulting funds from, Navire Pharmaceuticals. A member of his immediate family has equity in Array Pharmaceuticals and Mirati Pharmaceuticals, Inc.

